# Induction of Senescence During Postpartum Mammary Gland Involution supports tissue remodeling and promotes postpartum tumorigenesis

**DOI:** 10.1101/2024.12.29.630686

**Authors:** Aurelie Chiche, Elsa Charifou, Lamia Djoual, Shuoyang Wang, Laurianne Temime, Marielle Saclier, Shaoxiang Wang, Jeremy Chantrel, Han Li

**Affiliations:** Cellular plasticity in age-related pathologies, Dept. of Developmental & Stem Cell Biology, CNRS UMR 3738, Université Paris Cité Institut Pasteur, 25 rue du Dr Roux, Paris 75015, France; Sorbonne Université, Collège Doctoral, F-75005 Paris, France

**Keywords:** cellular senescence, tissue repair, mammary gland involution, postpartum breast cancer

## Abstract

Cellular senescence is an evolutionarily conserved stress response^1^, yet its roles during physiological processes remain underexplored^2,3^. Senescent cells are known to promote tissue repair^4,5^ and suppress tumorigenesis^6^, but their accumulation contributes to various pathological and physiological processes, including cancer and ageing^7–9^. However, it is currently unknown whether physiological senescence can be co-opted by oncogenic events to promote tumorigenesis. Postpartum mammary gland involution is a major tissue remodelling event in adulthood^10,11^, resembling the wound healing process, and is closely linked to postpartum breast cancer (PPBC)^12^ providing a compelling context to investigate this question. Here, we show that senescence is induced in alveolar luminal cells during involution in a p16-dependent manner. Reducing senescent cells hinders tissue remodeling and delays involution, underscoring their physiological importance. However, using a PPBC mouse model where the oncogenic event coincides with involution, we demonstrate that eliminating involution-associated senescent cells markedly extended the cancer latency. Mechanistically, we reveal that senescent cells enhance tumor-initiating cell plasticity in a paracrine manner, promoting tumor invasion and metastasis. Collectively, our findings uncover a dual role of senescence in mediating postpartum tissue remodeling and promoting tumorigenesis, highlighting a scenario where physiological senescence is hijacked to drive cancer progression. This work underscores that senescence might be a unifying mechanism linking tissue repair to tumorigenesis.

## INTRODUCTION

Cellular senescence is a stress-induced cell state characterized by stable cell cycle arrest, resistance to apoptosis, and a robust secretome known as senescence-associated secretory phenotype (SASP)^7,9^. However, senescence is not a uniform identity as previously viewed, it is now recognized as a highly heterogeneous phenotypes and pleiotropic functions that vary depending on cell type origin, inducers, and tissue context^1^. The stable cell cycle arrest is a potent cell-intrinsic tumour suppression mechanism ^6^, yet contributes to diminished regenerative capacity and tissue dysfunction during aging^13^. Extrinsically, senescent cells modulate extensively the microenvironment by secreting SASP factors and presenting cell-surface proteins^14,15^. The functional duality positions senescence as a key player in numerous physiological and pathological processes^7,9^, including embryonic development^16,17^, tissue repair^4,5^, fibrosis^18–20^, cancer^8^, and ageing^21^. Importantly, targeted elimination of senescent cells can delay the onset and improve many ageing-associated pathologies^22–24^. Despite extensive research into the pathological roles of senescence, its functional contributions to normal physiology remain poorly understood^2,3^. Furthermore, it remains unclear whether physiological senescence can be hijacked by oncogenic processes to drive tumorigenesis.

The postpartum period involve extensive physiological changes with lasting implications for women’s health^25,26^. Among these, the mammary gland undergoes dramatic remodeling during postpartum involution, a process that returns the gland to its pre-pregnant state after lactation (or directly after pregnancy if lactation does not occur)^10^. During this process, all the newly formed lobuloalveolar structures generated during pregnancy are removed via apoptosis and lysosomal-mediated cell death^27–32^. Concurrently, the gland undergoes extensive remodeling of the extracellular matrix, blood vessels, and adipocytes refilling, companied by a strong inflammatory response and immune infiltration^33–40^, to restore mammary epithelium integrity. Of note, mammary involution resembles a scarless wound-healing process as the mammary epithelium after involution can respond properly to the subsequent pregnancy and lactation repetitively throughout female’s reproductive life span.

While involution is critical for mammary gland homeostasis and the successful establishment of future lactation^11^, it also create a transient pro-tumorigenic microenvironment^12^ and is strongly implicated in postpartum breast cancer (PPBC), defined as breast cancer diagnosed within 5–10 years after childbirth^41–43^. PPBC is associated with higher metastasis rates and poorer survival compared to nulliparous women or those diagnosed during pregnancy, regardless of hormone receptor status^41–43^. Notably, increased maternal age positively correlates with a higher incidence of PPBC^44^. With the global trend of delayed childbirth, PPBC incidence and mortality rates are expected to rise significantly, emphasizing a critical unmet medical need. However, the mechanisms underlying the pro-tumorigenic features of involution remain not well understood.

Here, we provide evidence that certain alveolar luminal cells upregulate p16 to acquire senescence state during postpartum mammary gland involution. Loss-of-functions studies reveal that senescence is required for proper tissue remodelling during involution but, paradoxically, also promotes tumorigenesis and metastasis in a paracrine manner by enhancing tumour-initiating cells plasticity. Thus, our findings demonstrate that in a physiological context like mammary gland involution, senescence plays a dual role both mediating tissue remodeling and facilitating tumorigenesis and suggesting that senescence might serve as a common mechanism linking wound healing to cancer.

## RESULT

### Cellular senescence is transiently induced at the onset of the tissue remodeling phase of mammary gland (MG) involution

To investigate whether senescence occurs throughout MG postnatal development and the reproductive cycle, we performed senescence-associated beta-galactosidase (SA-βGal) staining on murine MGs collected at various developmental stages, including puberty (6 weeks), mature virgin (10 weeks), pregnancy (p13.5 and p18.5), lactation (lac10), and involution (Figure 1A). Notably, robust SA-βGal staining was observed in involuting MGs, while SA-βGal+ cells were scarcely detectable at other stages (Figure 1B). The involution process in rodents consists two morphologically distinct phases^10^. During the first 48 hours, programmed cell death occurs, while the alveolar lumen remains intact. This is followed by a phase of extensive tissue remodeling between 60 and 72 hours, characterized by alveolar regression and adipocyte repopulation^10^. To analyze the dynamics of SA-βGal+ cells during involution, we initiated synchronized involution by removing pups after 10 days of lactation and collected MGs at different time points: Inv1, Inv3, Inv7, Inv14, and Inv28 (representing the resting MG) (Figure 1A). Interestingly, SA-βGal+ cells became apparent starting at Inv3, with staining intensity peaking at Inv7 and gradually diminishing thereafter (Figure 1C, i-iv). No SA-βGal+ cells were detected in the resting MG following the completion of involution (Inv28, Figure 1C, v). Most SA-βGal staining was spatially restricted to the alveolar region compared to the ductal epithelium.

**Figure 1.**
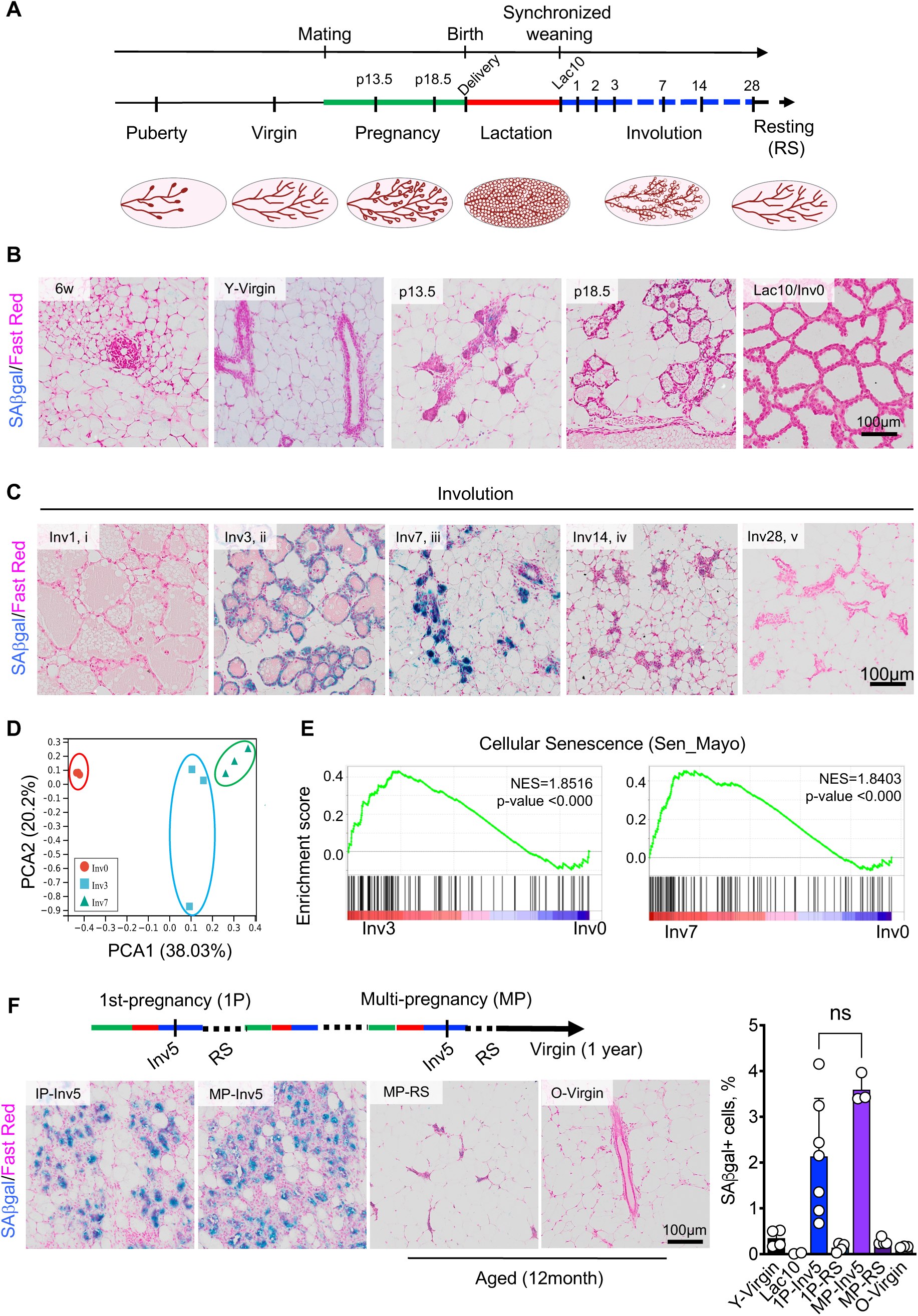
Transient induction of senescence during mammary gland involutio. **A.** Experimental design highlighting the time points of mammary glands collection at: puberty (6 weeks), adult virgin (10 weeks), pregnancy (p13.5, p18.5), lactation (10 days - Lac10/Inv0), involution after 1, 3, 7, 14 and 28 days/resting stage (Inv1, Inv2, Inv3, Inv7, Inv14, and Inv28/RS). **B-C.** SAβgal staining during MG developmental stages. Scale bar: 100μm. **D.** Principal Component Analysis (PCA) of transcriptomes from mammary glands collected at Inv0, Inv3, Inv7 samples (n=3 for each). **E.** GSEA enrichment plots. Reference gene set: SEN_MAYO. **F.** SAβgal staining of mammary gland during the 1^st^ involution (1P-Inv5), after at least 3 rounds of reproductive cycles (MP-Inv5 and MP-RS), or in old virgin females (12 months). Scale bar: 100μm (left panel). Quantification of SAβgal cells related to total number of cells at different timepoints of mammary gland stage (right panel), Y-Virgin n=4 ; Lac10 n=2 ; 1P-Inv5 n=7 ; 1P-RS n=4 ; MP-Inv5 n=3 ; MP-RS n=4 ; O-Virgin n=3. Data are represented as mean ± standard deviation.

To further validate our observations and characterize the senescence program associated with mammary gland involution, we conducted bulk RNA sequencing (RNA-seq) on MGs from Inv0, Inv3, and Inv7. Principal component analysis (PCA) revealed clear clustering of Inv0, Inv3, and Inv7 samples, indicating distinct biological differences across these stages (Figure 1D). Gene-set enrichment analysis (GSEA) demonstrated significant enrichment of senescence-associated genes^45^ in both Inv3 and Inv7 compared to Inv0 (Figure 1E). Heatmap depicting upregulated senescence-associated genes in Inv3 and Inv7 relative to Inv0 further highlighted this senescence signature (Supplementary Figure 1A). Interestingly, we identified numerous genes crucial for the involution process, including extracellular matrix (ECM) remodeling factors such as MMP2, MMP3, MMP14, and Serpine1^36,37^; cytokines and chemokines^46^ like CXCL12, CXCL10, CCL4, CXCL16, CCL3, CCL2, IL18, and CCL8; and growth factors^37,47^, including GDF15, FGF2/7, and IGFBP4/5. Additionally, hallmark gene sets frequently associated with senescence, such as interferon responses^48^, epithelial-mesenchymal transition (EMT)^49^, NF-κB signaling^50^, and the complement system, were significantly enriched in Inv3 and Inv7 (Supplementary Figure 1B). Collectively, these findings suggest that senescence is transiently induced during mammary gland involution, coinciding with the onset of tissue remodeling.

### Involution-associated senescence does not accumulate with age

Senescent cells accumulate during aging in various tissues^9^. A recent study has reported that aged mammary stem/progenitor cells acquire a gene signature associated with cellular senescence^51^. We questioned whether senescent cells might accumulate in the mammary gland upon multiple pregnancies (multiparity). To address this question, we compared SA-βGal^+^ cells across several groups, including involuting MGs following the first pregnancy (1P-Inv5, ∼3 months), resting mammary glands after one round of involution (1P-RS ∼4 months), involuting MGs following multiparous glands (MP-Inv5, ∼11 months, corresponding to approximately 35 years in humans), resting mammary glands post-MP (MP-RS ∼12 months, corresponding to approximately 40 years in humans), and glands from nulliparous mice (O-virgin ∼12 months) (Figure 1F). We observed a minor, non-significant increase in SA-βGal^+^ cells in involuting multiparous MGs (MP-Inv5) compared to 1P MGs (IP-Inv5) (Figure 1F). Furthermore, we detected negligible SA-βGal^+^ cells in mammary glands from one-year-old females, regardless of parity status. These findings suggest that the senescence program associated with involution does not accumulate or amplify with multiple pregnancies during reproductive aging.

### Alveolar luminal cells are the major senescent cell type

To identify which cell types undergo senescence during MG involution, we employed C_12_FDG, a fluorogenic substrate for β-galactosidase ^52^, alongside cell-specific markers. Stromal cells were identified as CD31^-^, CD45^-^, CD24^-^, α6^-^; basal cells as CD31^-^, CD45^-^, CD24^low^, α6^high^; and luminal cells as CD31^-^, CD45^-^, CD24^high^, α6^low^. Using flow cytometry, we found that approximately 6% of stromal cells, 15% of basal cells, and 52% of luminal cells were C_12_FDG^+^ at Inv5 compared to Inv0 (Supplementary Figure 2B). Although our antibody panel could not distinguish alveolar epithelial cells from ductal epithelial cells, histological staining indicated that most SA-βGal^+^ cells are alveolar epithelial cells. Next, we conducted co-staining of SA-βGal with the basal cell marker Keratin 5 (Krt5) and the luminal cell marker Keratin 8 (Krt8) at Inv3. Consistently, we found that the majority of SA-βGal^+^ cells were Krt8^+^ alveolar luminal cells (Figure 2A), which are negative for both the apoptotic marker cleaved Caspase-3 (CC3) and the proliferation marker Ki67 (Supplementary Figure 2C & 2D). Together, our findings indicate that alveolar luminal cells are the primary senescent cell type during mammary gland involution.

**Figure 2.**
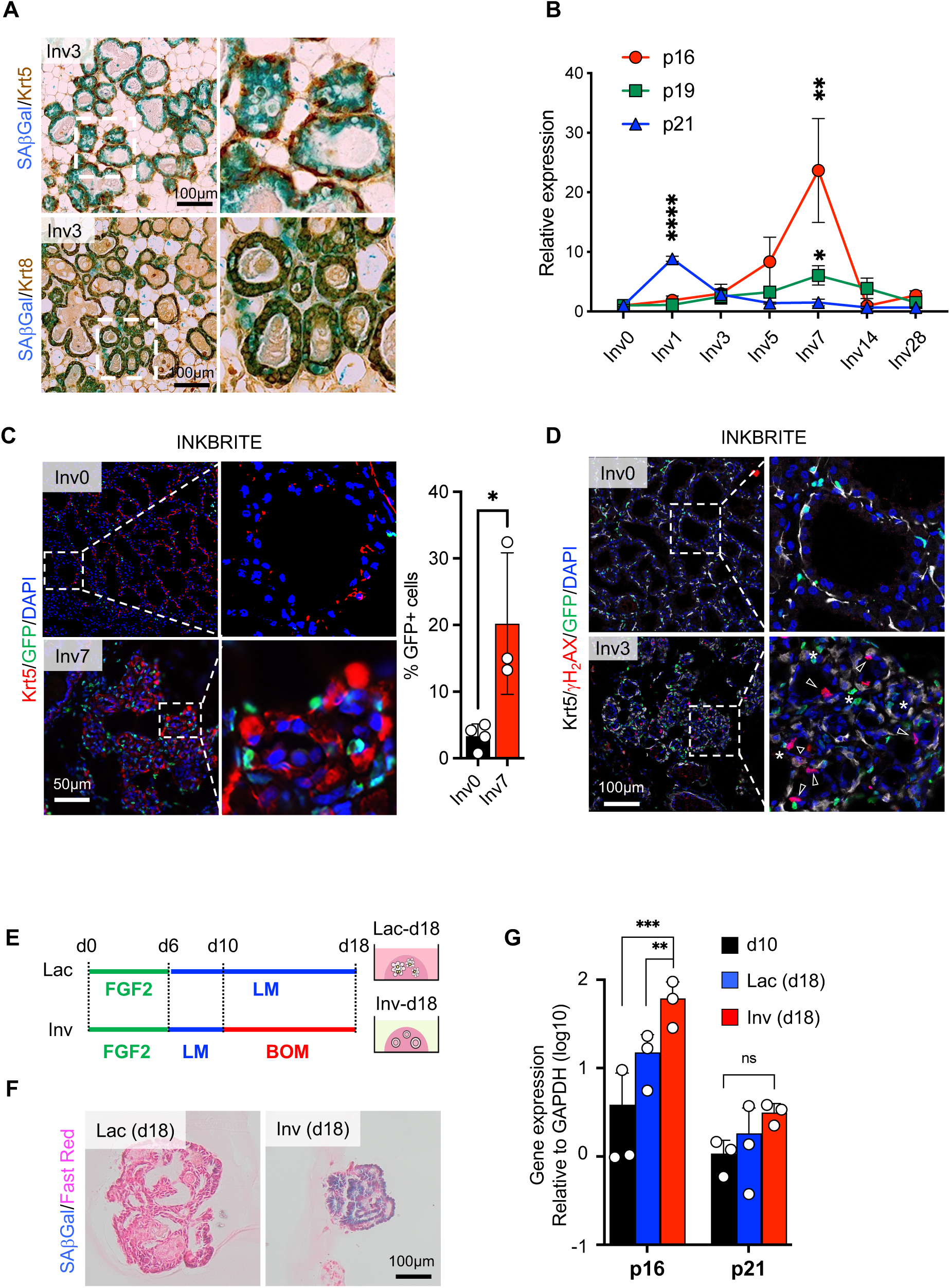
Alveolar luminal cells are the major senescent cell type. **A.** Representative pictures of co-staining SAβGal with Krt5 or Krt8 markers on WT mammary glands from Inv3 females (n=2). Scale bar: 100μm **B.** qPCR analysis of p16, p19 and p21 genes from mammary glands collected from Inv0 to Inv28 (n=4 for each timepoint). Data are represented as mean ± standard deviation relative to Inv0. Two-way ANOVAs were performed. p-values *<0.05, **<0.01, ****<0.0001. **C.** Representative pictures of co-staining of GFP with Krt5 on INKBRITE involuting mammary gland from Inv0 (n=4) and Inv7 (n=3). Scale bar: 50μm. Quantification represents p16^INKBRITE-GFP+^ cells. **D.** Representative pictures of co-staining of GFP with Krt5 and gH_2_AX on INKBRITE involuting mammary gland from Inv0 (n=4) and Inv3 (n=4). Scale bar: 100μm **E.** Experimental design of *ex vivo* culture of organoids. **F.** SAβGal staining of involuting organoids from involuting (BOM, d18) and lactation control (LM, d18) (n=3). Scale bar: 100μm **G.** qPCR analysis of p16 and p21 of organoids from involution-like and lactation-like process. n=3 biological replicates. Data are represented as mean ± standard deviation. One-way ANOVAs were performed. p-values **<0.01, ***<0.001.

### p16 upregulation, but not p21, is associated with mammary involution-induced senescence

Interestingly, our bulk RNAseq analysis revealed upregulation of *Cdkn2a*, *Rb1*, *Trp53*, and various Trp53 target genes^53^, but not *Cdkn1a*, at Inv3 and Inv7 compared to Inv0 (Supplementary Figure 2E & 2F). To validate these findings, we performed qPCR on RNA extracted whole MGs at different involution time points. Results showed significant upregulation of both *p16^INK4a^* and *p19^Arf^* at Inv7, while *p21* was transiently upregulated at Inv1 and returned to the basal levels by Inv3 (Figure 2B). We further investigated p16INK4a expression using a sensitive reporter mouse model, INKBRITE, where the p16^INK4a^ promoter drives expression of multiple copies of H2B-GFP^54^. At Inv0, few GFP^+^ cells were observed within alveolar structure marked by Krt5 or in the stromal compartment (Figure. 2C, upper panel). Importantly, at Inv7, a significantly higher proportion of GFP^+^ cells was detected, predominantly within the alveolar structures facing the lumen (Figure 2C, lower panel). Next, we examined whether involution-induced senescence was triggered by DNA damage. While we observed γH2AX+ cells at Inv3, predominantly in alveolar luminal cells, GFP^+^ and γH_2_AX^+^ cells were mutually exclusive (Figure 2D). These findings suggest that involution-induced senescence is linked to p16 upregulation, independent of p21, and occurs most likely independently of DNA damage.

We recently established a mammary organoid system that models mammary postnatal developmental stages including pregnancy, lactation, and involution *ex vivo*^55^. To test whether senescence is also induced during involution-like process in this organoid model, we performed whole-mount SAβGal staining on organoids cultured in either lactation-like medium (LM, Lac) or involution-like medium (BOM, Inv) (Figure 2E). Histological analysis revealed that involution-organoids were notably smaller than lactation-organoids, with significantly more SA-βGal^+^ cells in the involution-organoids (Figure 2F & Supplementary Figure 2G). Consistently, qPCR analysis confirmed significant upregulation of p16 in involution-like organoids, with no significant change in p21 expression (Figure 2G). Thus, the organoid model supports the *in vivo* finding that p16, but not p21, is associated with senescence during involution, and organoid system can serve as a simplified *ex vivo* model to study involution-associated senescence.

### Systemic lactogenic hormones prevent senescence induction during involution

Next, we investigated upstream signals regulating senescence induction during involution. Organoid experiments indicated that prolactin withdrawal triggers senescence (Figure 2E-G). To assess if systemic lactogenic signals are important for senescence induction *in vivo*, we sealed a set of teats of lactating mice to initiate local involution while maintaining systemic lactogenic hormone levels (Figure 3A & Supplementary Figure 3A). It is known that involution-associated programmed cell death (PCD) is triggered locally^28^, and consistent with previous findings, we observed distinct morphological differences between involuting and sealed MGs at the onset of irreversible involution (day 3, Figure 3B & Supplementary Figure 3C & 3D). The sealed MGs displayed intact alveolar structures without tissue remodeling, unlike the involuting MGs. Importantly, no SA-βGal^+^ cells or p16 upregulation were detected in sealed MGs compared to involuting MGs at Inv3 (Figure 3B, Supplementary Figure 3B). These results suggest that systemic lactogenic hormones withdrawal upon weaning is required for senescence induction during involution.

**Figure 3.**
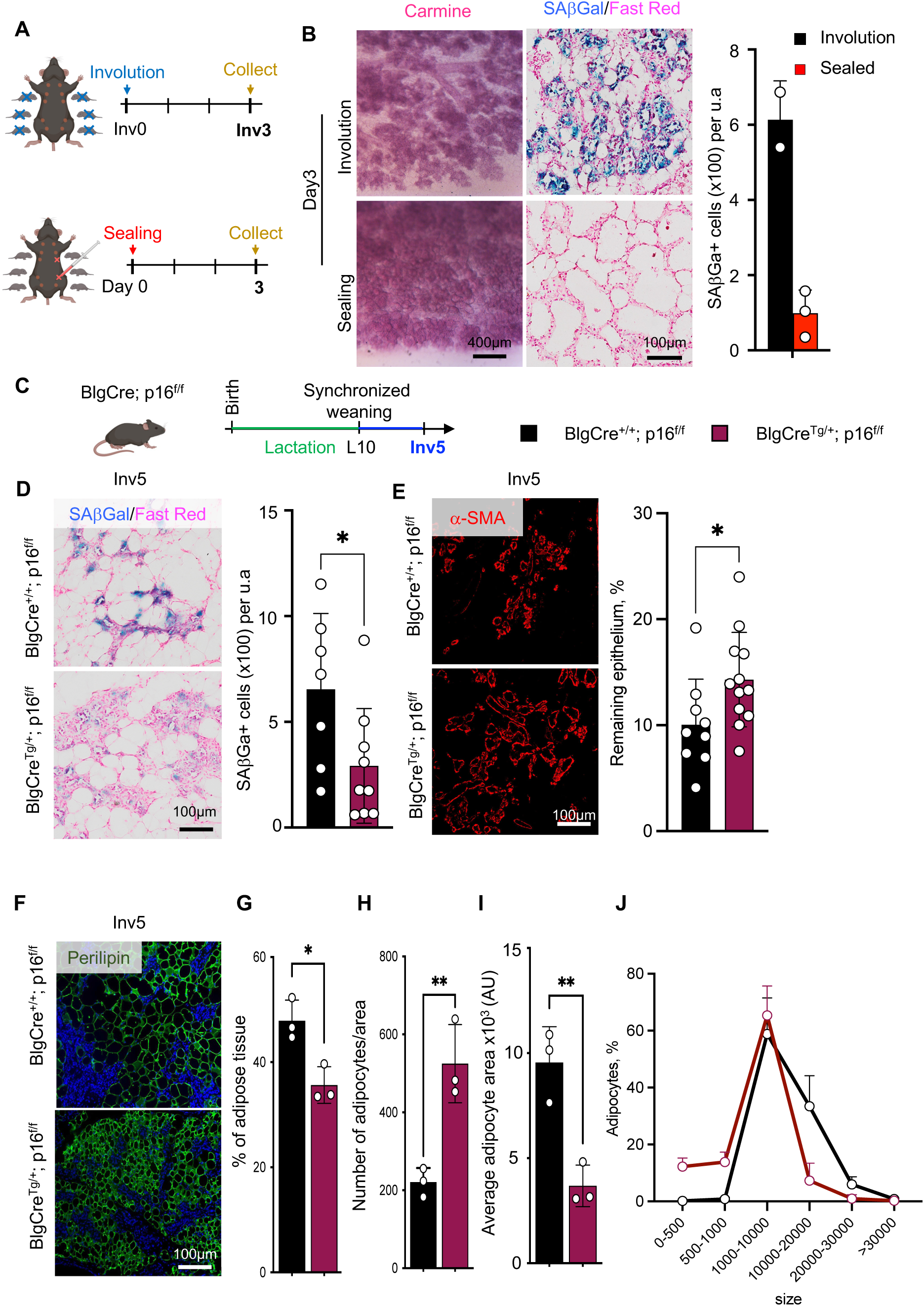
Impaired senescence delays mammary gland involution. **A.** Experimental design of involuting (blue) or teat sealed (red) mammary glands, while other mammary glands are maintained in lactation. Mammary glands were collected 3 days after sealing and compared with mammary glands from mice undergoing normal involution. **B.** Representative pictures of whole mount carmine and SAβGal staining from involuting or sealed mammary glands at day 3, on cryosections, counterstained with FastRed. Scale bar: 100μm. Automatic quantifications of SAβGal staining at day 3, from involuting (black, n=2) or sealed (red, n=3) mammary glands. Data are represented as mean ± standard deviation. **C.** Experimental design of experiment with BlgCre; p16^f/f^ mice. **D.** Representative pictures of SAβGal staining (left panel) from Blg^+/+^; p16^f/f^ or Blg^Tg/+^; p16^f/f^ mammary glands at Inv5, on cryosections, counterstained with FastRed. Scale bar: 100μm. Automatic quantification of SAβGal staining (right panel), Blg^+/+^; p16^f/f^ n=7 ; Blg^Tg/+^; p16^f/f^ n=9. Data are represented as mean ± standard deviation. Unpaired t-test was performed. p-values *<0.05. **E.** Representative pictures of α-SMA staining (left panel) from Blg^+/+^; p16^f/f^ or Blg^Tg/+^; p16^f/f^ mammary glands at Inv5. Scale bar: 100μm. Quantification of remaining epithelium (right panel), Blg^+/+^; p16^f/f^ n=9 ; Blg^Tg/+^; p16^f/f^ n=12. Data are represented as mean ± standard deviation. Unpaired t-test was performed. p-values *<0.05. **F.** Representative pictures of perilipin staining from Blg^+/+^; p16^f/f^ or Blg^Tg/+^; p16^f/f^ mammary glands at Inv5. Scale bar: 100μm. **G-I.** Quantification of adipose tissue proportion (H), number of adipocytes per unit area (I), and average of adipocyte area per unit area (J), n=3 mice per group. Data are represented as mean ± standard deviation. Unpaired t-test was performed. p-values *<0.05, **<0.01. **J.** Distribution of adipocytes size from Blg^+/+^; p16^f/f^ or Blg^Tg/+^; p16^f/f^ mammary glands at Inv5, n=3 mice per group. Data are represented as mean ± standard deviation.

### p16-dependent senescence induction ensures involution dynamic

To determine the functional relevance of senescence in mammary involution, we used both genetic and pharmaceutical approaches to perturb the senescence program. Firstly, we generated BlgCe^Tg/+^;*p16^f/f^*mice, an alveolar luminal cell-specific p16 knockout model, to assess p16’s role in senescence during involution. In BlgCe^Tg/+^;*p16^f/f^* mice, we observed a delayed onset of tissue remodeling compared to wild-type controls (Supplementary Figure 3E). Importantly, BlgCe^Tg/+^;*p16^f/f^*MGs exhibited significantly fewer SAβGal^+^ cells at Inv5 compared to littermate controls (Figure 3D), though a minor, non-significant reduction in SAβGal^+^ cells was also observed at Inv4 (Supplementary Figure E). Hence, these findings demonstrate that alveolar luminal cell senescence during involution is p16 dependent.

Next, we evaluated the functional consequence of impaired senescence induction during MG involution, using alveolar size and adipocyte area as metrics to evaluate the involution kinetic. Interestingly, at Inv5, BlgCe^Tg/+^;*p16^f/f^*MGs retained more epithelial structures and exhibited a reduced adipocyte area compared to controls (Figure 3E-G). Additionally, BlgCe^Tg/+^;*p16^f/f^* MGs contained much more, but markedly smaller adipocytes than controls (Figure 3H-J), suggesting a link between reduced senescence and impaired adipocyte re-differentiation and hypotrophy. Collectively, these findings demonstrate that p16-dependent senescence is crucial for proper mammary gland involution.

### ABT-263 treatment delays mammary gland involution

Next, we used ABT-263^23,56^, a BH3 mimetic targeting anti-apoptotic Bcl-2 family proteins, to deplete senescent cells during involution. Mice were treated with ABT-263 daily for 3 days prior, and MGs were harvested at Inv3, Inv4, and Inv5 (Fig. 4A). Notably, ABT-263 treatment led to a significant increase in apoptotic (CC3+) cells and a decrease in SA-βGal^+^ cells, suggesting apoptosis was induced in senescent cells (Figure 4B & 4C). Notably, ABT-263 did not affect STAT3 activation, a key mediator of PCD during mammary involution^57^ (Supplementary Figure 4A). Consistent with the BlgCre^Tg/+^; *p16^f/f^* mice, ABT-263-treated MGs exhibited delayed alveolar regression and reduced adipocyte refilling at Inv4 and Inv5 compared to controls (Figure 4D & 4E). However, unlike the BlgCre^Tg/+^; *p16^f/f^* model, we observed no change in adipocyte numbers, except at Inv3 (Supplementary Figure 4C), indicating differences between senescence inhibition and senescent cell clearance. These findings indicate that ABT-263 effectively removes senescent cells and delays the involution process. Taken together, both genetic and pharmaceutical approaches demonstrated that senescence induction is important for proper MG involution.

**Figure 4.**
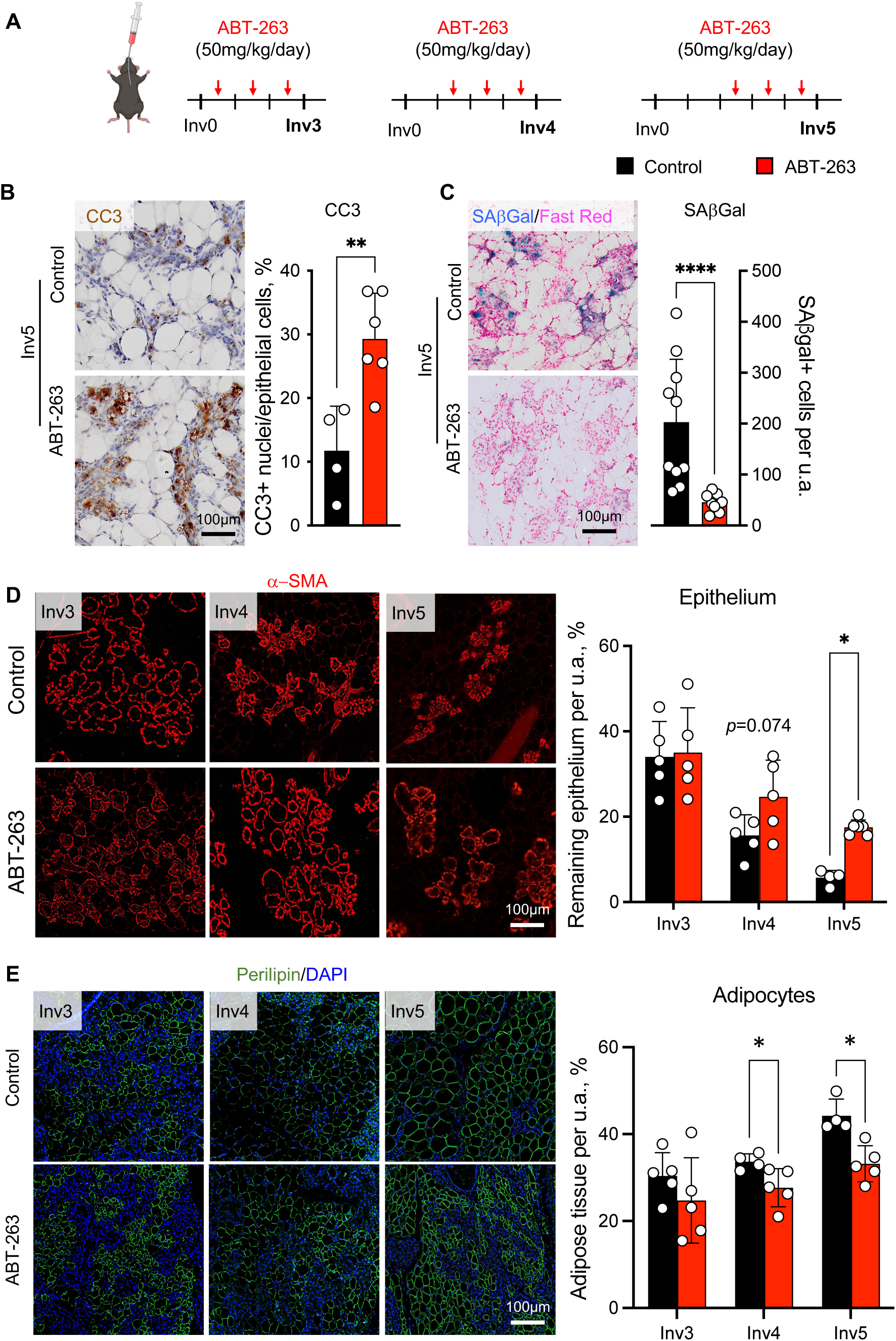
ABT-263 treatment delays mammary gland involution. **A.** Experimental design of *in vivo* treatment with ABT-263 at 50mg.kg^-1^.day^-1^ or vehicle. Mice were force-fed every day for 3 days prior to mammary gland collection at Inv3, Inv4 or Inv5. **B-C.** Representative pictures of mammary gland sections from control or ABT-263 treated mice at Inv5, respectively stained for cleaved-caspase 3 (CC3) (B) and SAβGal on cryosection (C). Scale bar: 100μm. Quantifications of CC3 (B) and SAβGal staining (C). CC3 staining: Control n=4; ABT-263 n=6. SAβGal staining: Control n=10; ABT-263 n=8. Data are represented as mean ± standard deviation. Unpaired t-tests were performed to analyze the effect of ABT-263 treatment respectively on the number of CC3^+^ or SAβGal^+^ cells. p-values **<0.01, ****<0.0001. **D-E.** Representative pictures of mammary gland sections from control or ABT-263 treated mice at Inv3 to Inv5, respectively stained for α-SMA (upper panels), or Perilipin (lower panels). Scale bar: 100μm. Quantifications (right panels) for remaining epithelium and proportion of adipose tissue in percentage. α-SMA staining: Inv3/Inv4 n=5 for each group; Inv5 Control n=4, Inv5 ABT-263 n=6. Perilipin staining: Inv3 n=5 for each group; Inv4/Inv5 Control n=4, Inv4/Inv5 ABT-263 n=5. Data are represented as mean ± standard deviation. Unpaired t-tests were performed. p-values *<0.05.

### Senescent cells recruit macrophages during involution

To gain insights on the role of senescence in involution, we further analyzed our bulk RNA-seq datasets. GSEA analysis revealed enriched biological processes related to immune cell chemotaxis and migration (Figure 5A and Figure 5B). We took advantage of the involuting organoid system to determine the expression of relevant genes (Figure 5C). The expression of *Csn2*, important for milk production, was significantly induced by prolactin and reduced after prolactin withdrawal^55^(Supplementary Figure 5A). Consistently, p16 expression was significantly upregulated post prolactin withdrawal (Supplementary Figure 5B). Interestingly, we found that chemokines critical for monocyte/macrophage recruitment and ECM modifiers important for tissue remodeling were upregulated in involuting organoids compared to lactational controls (Figure 5D, Supplementary Figure 5B), suggesting that senescent epithelial cells are at least one of the sources producing these factors.

**Figure 5.**
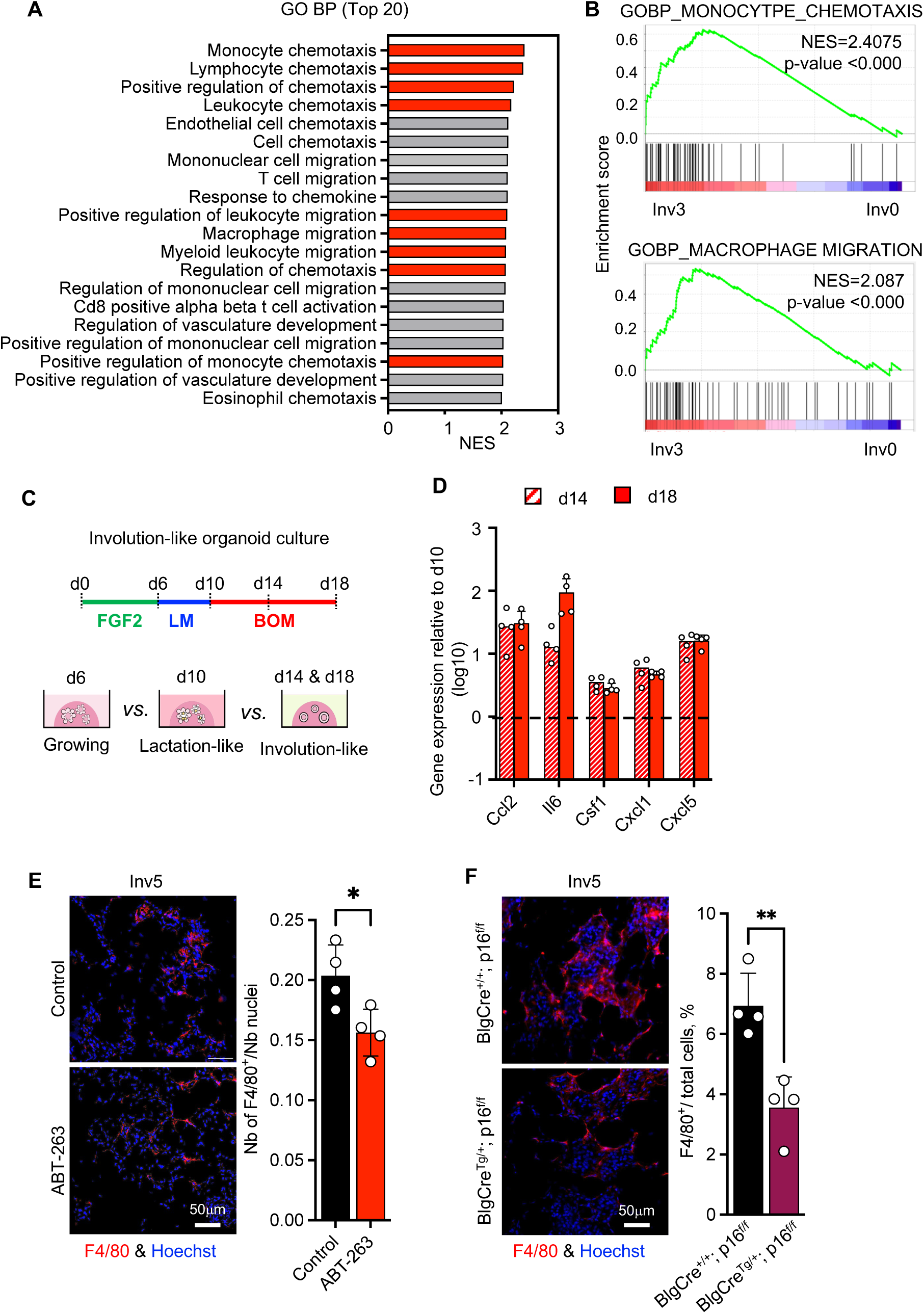
Involution-induced senescence is required for macrophage recruitment. **A.** GSEA analysis of GO-Biological Processes, top 20 enriched pathways. **B.** GSEA enrichment plots. Reference gene set: GOBP_MONOCYTE_CHEMOTAXIS (upper panel) and GOBP_MACROPAGE_MIGRATION (lower panel). **C.** Design of the organoids culture experiment. d0 to d6: FGF_2_ treatment to induce growth of organoids. d6 to d10: addition of lactogenic medium (LM) to mimic lactation. d10 to d18: Basic organoid medium (BOM) to mimic involution. **D.** qPCR analysis of Ccl2, IL6, Csf1, Cxcl1, Cxcl5 genes from mammary organoids collected at d14 and d18 relative to d10. n=4 for each timepoint. Data are represented as mean ± standard deviation. **E.** Representative pictures of mammary gland sections from control or ABT-263 treated mice at Inv5, stained for F4/80 and its quantification. n=4 mice per group. Data are represented as mean ± standard deviation. Mann-Whitney test was performed. p-values *<0.05. Scale bar: 50μm **F.** Representative pictures of mammary gland sections from Blg^+/+^; p16^f/f^ or Blg^Tg/+^; p16^f/f^ mice at Inv5, stained for F4/80 and its quantification. n=4 mice per group. Data are represented as mean ± standard deviation. Unpaired t-test was performed. p-values **<0.01. Scale bar: 50μm.

Macrophages are known to infiltrate involuting MG at the onset of tissue remodeling phase, a process essential for adipocyte repopulation and involution progression^58,59^. Considering the intimate relationship between macrophage and senescent cells^60^, we hypothesized that senescent epithelial cells recruit macrophages during involution. Supporting this, we found significantly fewer CD45^+^ immune cells and macrophages (CD64^+^ or F4/80^+^) in ABT-263-treated involuting MGs compared to controls (Figure 5E, Supplementary Figure 5C-E). Consistently, BlgCre^Tg/+^; *p16^f/f^* involuting MGs showed fewer F4/80^+^ macrophages than controls (Figure 5F). These data suggest that senescent epithelial cells play a key role in recruiting macrophages during involution to ensure proper involution dynamic.

### Involution associated senescence promotes postpartum tumorigenesis

Next, we set to explore the potential implication of involution-associated senescence in breast cancer. Women diagnosed with breast cancer within five to ten years postpartum exhibit a worse prognosis compared to age-matched nulliparous women or those diagnosed during pregnancy independent of hormone receptor status^12,41^. It has been proposed that the involuting mammary gland microenvironment promotes breast cancer metastasis^44,61,62^. Given senescent cells can extensively rewire the microenvironment to influence cancer progression in a context-dependent manner ^8,63–65^, we investigated whether senescence associated with mammary gland involution contributes to a pro-tumorigenic microenvironment.

To test this hypothesis, we utilized the MMTV-neu mouse model (Figure 6A). Consistent with existing literature^62^, we observed that a single round of pregnancy and subsequent involution significantly reduced tumor latency (Figure 6B). Surprisingly, repeated rounds of pregnancy mitigated the impact of parity on tumorigenesis (Figure 6B). Previous studies suggest that concurrent pregnancy delays the involution process^66^. To examine whether concurrent pregnancy affects involution-associated senescence, we mated MMTV-neu mice and dispatched them into two groups randomly. We removed the male from the control group after confirming pregnancy (control group), while left the male with the other group of pregnant females (concurrent pregnancy group).

**Figure 6.**
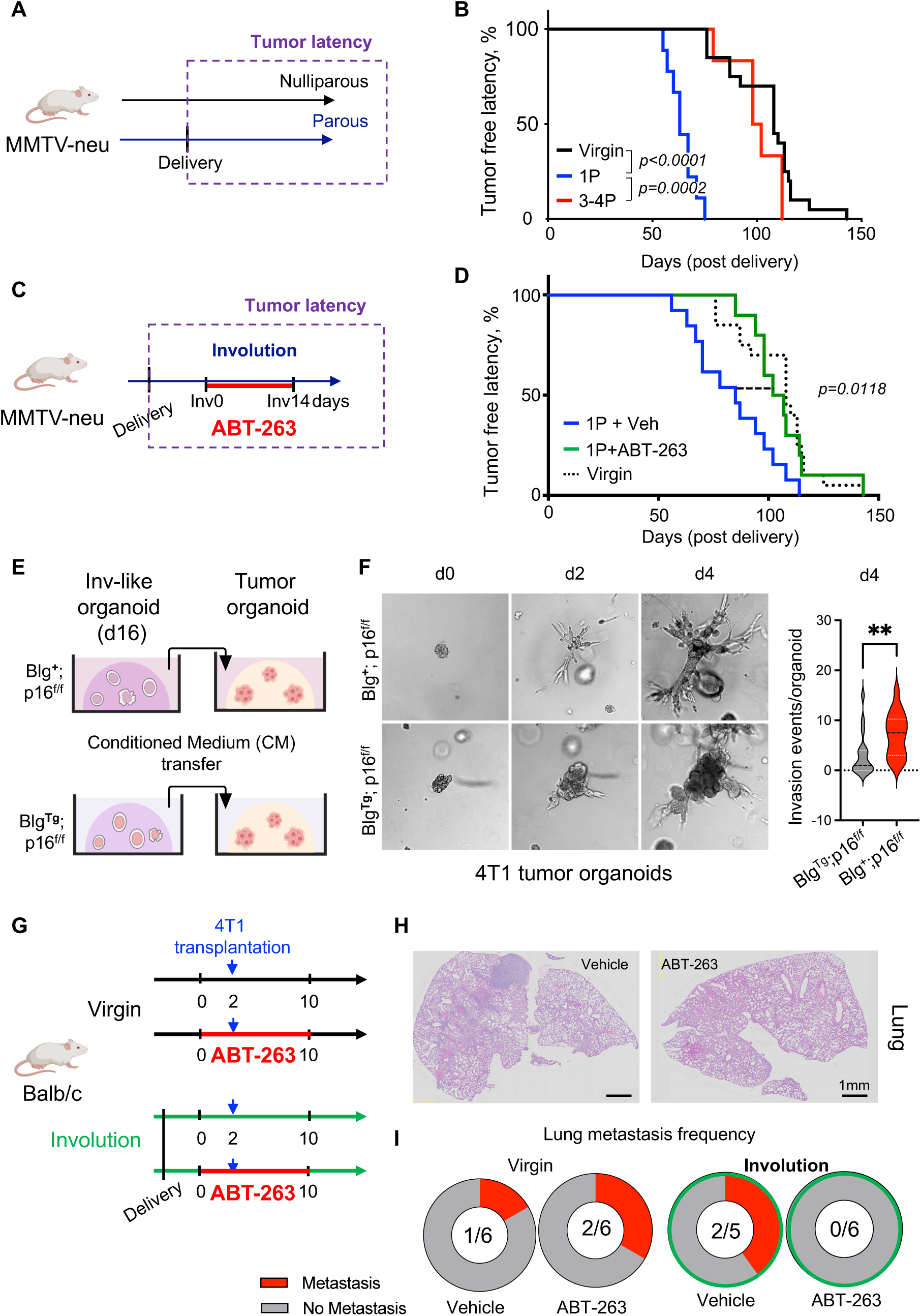
Involution-induced senescence promotes postpartum tumorigenesis. **A.** Experimental design of *in vivo* experiment with MMTV-neu mice. Mice were mated at the age of 10-12 weeks and allowed to go to lactation and natural involution, 1 to 3-4 times (called 1P or 3-4P for the number of pregnancies). Part of the littermate were maintained virgin to compare the tumor free latency. **B.** Tumor free latency curve comparing Virgin vs. 1 pregnancy (1P) vs. 3-4 pregnancies (3-4P) from the first date of delivery used as starting point for the virgin littermate. n=20 for virgin; n=9 for 1P; n=6 for 3-4P. Curve comparison Gehan-Breslow-Wilcoxon test was performed. p-value ****<0.0001 (Virgin vs. 1P); p-value=0.0002 (3-4P vs 1P). **C.** Experimental design of *in vivo* experiment with MMTV-neu mice treated with ABT-263 at 50mg.kg^-1^.day^-1^ or vehicle. Mice were force-fed every day for 2 weeks following synchronized involution and tumor development was monitored every week. **D.** Tumor free latency curve comparing 1 pregnancy + Vehicle (1P +Veh) vs. 1 pregnancy + ABT-263 (1P +ABT-263) from the date of delivery used as starting point. n=13 for 1P +Veh; n=10 for 1P +ABT-263. Curve comparison (Gehan-Breslow-Wilcoxon test) was performed. p-value *<0.0118. **E.** Experimental design of *in vitro* experiment. After collection, conditional media from Blg+; p16f/f or BlgTg; p16f/f was used to cultivate organoids isolated from 4T1 tumor (transplantation in virgin balb/c females). **F.** Representative brightfield pictures of tumor organoids growing from d0 to d4 (right panel). Quantification of invasion event per organoid at d4 (left panel), n=20 tumor organoids per condition. Unpaired t-tests were performed. p-values **<0.01. **G.** Experimental design of *in vivo* experiment for 4T1 tumor cells transplantation in Balb/c mice treated with ABT-263 at 50mg.kg^-1^.day^-1^ or vehicle. Mice were injected when virgin or at day 2 of involution. **H.** Representative pictures of lung sections from control or ABT-263 treated mice 4 weeks after transplantation, stained for hematoxylin and eosin. Scale bar: 1mm. **I.** Diagram of lung metastasis frequency in control or ABT-263 treated mice 4 weeks after transplantation. n=6 for Virgin +Veh; n=6 for Virgin +ABT-263; n=5 for Inv2 +Veh; n=6 for Inv2 +ABT-263.

Upon delivery, all females from both groups established lactation successfully, and we confirmed that females kept with the males (concurrent pregnancy) conceived again during lactation. We then removed pups from both groups to initiate synchronized involution and collected mammary glands at Inv3 and Inv7 (Supplementary Figure 6A). Histological analysis and STAT3 activation confirmed a delayed involution process in the concurrently pregnant females (Supplementary Figure 6B & 6C)^66^. Notably, we observed a significant reduction in SA-βGal^+^ cells in the involuting mammary glands of concurrently pregnant females (Supplementary Figure 6D & 6E). Our animal records further indicated that the multiparous MMTV-neu mice remained pregnant during involution. Thus, we hypothesized that bypassing involution-associated senescence might underlie the diminished impact of parity on tumorigenesis. To test this hypothesis, we treated MMTV-neu mice with ABT-263 for two weeks at the onset of involution (Figure 6C). Importantly, ABT-263 treatment significantly delayed tumor latency in parous mice compared to controls (Figure 6D), supporting the notion that involution-associated senescence promotes mammary tumorigenesis.

### Involution-associated senescence promotes tumor plasticity via SASP

Next, we sought to elucidate the mechanisms by which involution-induced senescence promotes tumorigenesis. Previously, we showed that senescence promotes cellular plasticity in the context of cellular reprogramming^67,68^, and the activation of Rank signaling induces senescence to promote mammary tumorigenesis^69^. Therefore, we investigated whether senescence promotes tumor plasticity by establishing a conditioned medium (CM)-based tumor organoid system. Specifically, we generated involuting organoids from both control and BlgCre^Tg/+^;*p16^f/f^* mice, collecting CM every two days for three times after day 14 (day 14, day 16, and day 18) to culture 4T1 tumor organoids derived from virgin Balb/c hosts (Figure 6E). Notably, tumor organoids cultured with CM from BlgCre^Tg/+^;*p16^f/f^*mice exhibited significantly fewer invasive events than those cultured with control CM (Figure 6F). These findings suggest that involution-associated senescence promotes tumor invasion in a paracrine manner.

### Involution associated senescence promotes mammary tumor metastasis in the lung

Finally, to investigate whether involution associated senescence could promote the tumor metastatic potential *in vivo*, we adapted a postpartum breast cancer mouse model focused on the tumor-promoting effects of involution^62^. We injected 4T1 cells into both virgin and involuting (Inv2) Balb/c mice, treating half of the groups with ABT-263 as described above (Figure 6G). Importantly, we found that ABT-263 treatment significantly suppressed lung metastasis (Figure 6H & 6I). Notably, no difference was observed in the virgin groups, indicating that ABT-263 specifically targets senescent cells in the involuting microenvironment without affecting tumor cells directly (Figure 6I). Taken together, these data support our hypothesis that involution associated physiological senescence enhances the plasticity of tumor-initiating cells to facilitate tumor invasion and metastasis.

Collectively, our study uncovers a pivotal dual role for senescence during postpartum mammary gland involution, demonstrating its importance in both physiological tissue remodeling and tumorigenesis.

## DISCUSSION

Here, we report that senescence is transiently induced during postpartum mammary gland involution, a physiological process critical for restoring mammary gland homeostasis after pregnancy. We identify that alveolar luminal cells are the primary cell type undergoing p16-dependent senescence, which is likely driven by systemic hormone withdrawal. Loss-of-function experiments demonstrate that senescence facilitates tissue remodeling processes to ensure proper involution. Unexpectedly, we also find that involution-associated senescence promotes PPBC progression by enhancing tumor-initiating cell plasticity through paracrine signaling. These findings reveal a previously unappreciated dual role for senescence in tissue repair and tumorigenesis, demonstrating how physiological senescence can be exploited by oncogenic events. This highlights the potential for targeting involution-associated senescence as a novel therapeutic strategy to mitigate postpartum breast cancer progression.

Involution-associated senescence shares notable parallels with senescence in decidual stromal cells of the cycling human endometrium^70^, which is required for successful implantation^71^. Both processes are triggered by fluctuation in female sex hormones, mediated by p53 and p16 upregulation, and play important roles in scarless tissue remodeling. However, their hormonal regulation differs. Brighton et al. showed that progesterone induces differentiation and senescence in human primary endometrial stromal cells during decidualization^70^, while in mammary gland, prolactin withdraw is required for senescence induction (Figure 2F, 2G and 3B). This context-dependent role of sex hormones underscores their importance as potential triggers of senescence. Interestingly, senescence induction in alveolar luminal cells appears to be DNA damage-independent (Figure 2D), consistent with development senescence^16^. Future work will be needed to delineate the molecular pathways that trigger mammary alveolar luminal cell senescence.

A recent study has reported that aged mammary gland stem cells exhibited senescence-like gene signature^51^. Curiously, our study showed that involution-associated senescent cells do not accumulate with age, which suggests that involution-associated senescence is a highly controlled and programmed process, distinct from the stochastic nature of ageing-associated senescence. In the future, it could be interesting to know whether senescent cells accumulate in the uterus during the premenopausal period.

Our RNA-seq analysis revealed that involution-associated senescence is linked to upregulation of *Cdkn2a* and *Trp53*. Notably, *Trp53^-/-^* and *p19Arf^-/-^* mice exhibit delayed mammary involution, highlighting the importance of these pathways in this process^72^. Interestingly, we did not observe a reduction in senescence burden or altered involution dynamics in *Cdkn2a^-/-^* mice (data not shown). However, acute deletion of p16 in alveolar luminal cells partially prevented senescence induction and impaired the involution process (Figure 3). These results suggest that potential compensatory mechanisms may obscure phenotypes in germ-line knockout models. It remains plausible that *p19^Arf^* or other members of the INK4 family also play critical roles in this process, warranting further investigation into their contributions.

To explore the functional relevance of senescence induction, we used both a genetically modified mouse model (BlgCre; *p16^f/f^*) and the senolytic compound ABT-263, which induces apoptosis in senescent cells. Both approaches significantly reduced senescent cell populations and delayed involution, notable differences were observed between the two strategies. In BlgCre; *p16^f/f^* mice, reduction in senescent cells and delayed involution became evident only at InvD5. In contrast, ABT-263-treated mice exhibited a reduction in senescence induction as early as InvD3, but this did not affect alveolar regression or adipocyte refilling until InvD4. These differences may be explained by the distinct mechanisms of action of the two approaches: the genetic model prevents only p16-dependent senescence, while ABT-263 actively removes all types of senescent cells through apoptosis. Intriguingly, we observed a significant increase in adipocyte numbers exclusively in BlgCre; *p16^f/f^* mice, suggesting that p16^+^ senescent cells might specifically impact adipogenesis.

Developmental and programmed senescence, such as those observed during embryonic development^16,17,73^, the female reproductive cycle^70,71^, and postpartum involution, including mammary gland and uterius^74^, occur within immune-privileged microenvironments. However, it remains unclear whether senescence actively establishes this immunosuppressive state. Our GSEA analysis revealed enrichment of pathways regulating immune cell chemotaxis, leading us to propose that senescent epithelial cells recruit macrophages during the irreversible phase of involution. Further characterization of the immune landscape will provide critical insights into the specific role of senescence in establishing immune suppression during postpartum tissue remodeling. Additionally, while both natural killer (NK) cells^71^ and macrophages^74^ are known to eliminate senescent cells in physiological contexts, it remains unknown which immune cells are required for regulating the fate of involution-associated senescence. Understanding these interactions will be pivotal in unraveling the immune-modulatory roles of senescence in postpartum tissue remodeling, which will have direct implications in tumorigenesis.

Finally, we demonstrated that physiological senescence program can be exploited by oncogenic process. In this study, we specifically focused on paracrine senescence in promoting tumor-initiating cells plasticity. Given the extensive body of research highlighting the distinctive features of involution^41–43^, it would be important to investigate whether and how senescence contributes to the establishment of a pro-tumorigenic microenvironment. Recent studies highlight shared immunosuppressive mechanisms between pregnancy and oncogenesis^75^, and senescent cancer-associated fibroblasts (CAFs) have been shown to mediate immune suppression to facilitate cancer progression^63,76^. These findings suggest potential parallels between the roles of senescence in physiological tissue remodeling and tumorigenesis, warranting further investigation.

## ACKNOWLEDGEMENTS

We are grateful to the Central Animal Facility, Cytometry Platform, and Bioinformatics and Biostatistics Hub of Institut Pasteur. This work was funded by Institut Pasteur, Centre National pour la Recherche Scientific and the Agence Nationale de la Recherche (Laboratoire d’Excellence Revive, Investissement d’Avenir; ANR-10-LABX-73), and Foundation ARC (PJA 20181208231). Work in the HL laboratory was also funded by Agence Nationale de la Recherche (ANR-16-CE13-0017, ANR-21-CE13-0006-01, ANR-22-CE16-0015-03), and AFMTELETHON (22403). Marielle Saclier is recipient of Revive postdoc fellowship, Elsa Charifou and Jeremy Chantrel are recipient of FRM 4^th^ PhD fellowships, Jeremy Chantrel’ PhD is funded by DIM Longévité et vieillissement PhD fellowship. Shuoyang Wang and Laurianne Temime were awarded Institut Pasteur Cancer Initiative M2 fellowship.

## AUTHOR CONTRIBUTIONS

A.C, E.C, L.D performed most of the experimental work, contributed to most of experimental design, data analysis, and discussions. SY. W, L.T, and M.S made critical experimental contributions. SX. W, and J.C performed bioinformatic analysis. H.L. supervised the study, designed the experiments and interpreted the data together with A.C, and wrote the manuscript. All authors discussed the results and commented on the manuscript. The authors declare no competing financial interests with this paper.

## DATA AVAILABILITY

The bulk RNA sequencing data have been deposited at Gene Expression Omnibus (GEO, access number GSE285005) and will be publicly available as of the date of publication. Source data supporting the findings of this study are provided with this paper. Additional data that support the findings of this study are available from the corresponding author upon reasonable request.

